# Influence of mimicry on extinction risk in Aculeata: a theoretical approach

**DOI:** 10.1101/2022.10.21.513153

**Authors:** Maxime Boutin, Manon Costa, Colin Fontaine, Adrien Perrard, Violaine Llaurens

## Abstract

Positive ecological interactions can play a role in community structure and species co-existence. A well-documented case of mutualistic interaction is Mullerian mimicry, the convergence of colour pattern in defended species living in sympatry. By reducing predation pressure, Mullerian mimicry may limit local extinction risks of defended species, but this positive effect can be weakened by undefended mimics (Batesian mimicry). While mimicry was well-studied in neotropical butterflies, it remains surprisingly poorly studied in wasps and bees (Hymenoptera: Aculeata). However, only females are defended in Aculeata and this female-limited defence may modulate the effect of Mullerian mimicry on extinction risks. Here, we focus on the effect of Mullerian mimicry on extinction risk in Aculeata, using a population dynamics model for two species. We show that Mullerian mimicry has a positive effect on species co-existence, but this effect depends on the sex-ratio. We found that the probability of extinction increases as the proportion of undefended males increases in the population, however co-existence still occurs if females are sufficiently abundant or noxious. Furthermore, we detected a destabilising effect of dual sex-limited mimicry (when each sex resembles a different model) on species co-existence. In a context of massive population decline caused by anthropic activities, our findings highlight the potential importance of Mullerian mimicry as an overlooked mechanism linked to extinction risk in wasp and bee species.

## Introduction

The assemblage of species within habitat is strongly driven by historical factors and abiotic constraints. Yet, community structure and co-existence of species are also likely to be shaped by ecological interactions, either antagonistic (*e*.*g*., competition, predation, parasitism) or facilitative interactions such as commensalism or mutualism (Holt, 2013) influencing both species colonization and extinction risk. The impact of antagonistic interactions on species co-existence and extinction have been extensively documented (Bruno et al., 2003), in particular the role of competition (Chesson, 2000; Raup, 1994). However, the impact of facilitative interactions on the composition of communities, and its underlying mechanisms, are much less studied. Such facilitation can involve improvement of physical environment (Bertness & Callaway, 1994), plant-pollinator interaction (Moeller, 2004) or associative defences against herbivores (Hay, 1986). Facilitative interaction between species plays a major role for species co-existence in stressful environments (Kéfi et al., 2008), and may prevail over the effect of competition (Gross, 2008). Positive ecological interactions have been shown to strongly affect species co-existence (Bastolla et al., 2009; Bertness & Callaway, 1994; Bronstein, 1994). In turn, mutualistic interactions may contribute to co-extinction dynamics, following the loss of a partner species. In the current context of biodiversity loss, studying the impact of mutualistic interactions on species extinction risk is thus especially relevant because co-extinction could be a major cause of species loss (Dunn et al., 2009; Koh et al., 2004).

Mullerian mimicry, whereby multiple defended species living in sympatry display similar conspicuous colour patterns, reducing individual predation risk (Müller, 1879), is a well-documented case of mutualistic interaction. This ecological interaction drives the convergence of warning patterns in defended species living in sympatry: local predators indeed learn the association between the conspicuous colour pattern of a prey and its defence (Rowland et al., 2007), therefore reducing predation rate on individuals sharing the same pattern (*i*.*e*., belonging to the same mimicry ring). This reduction on predation risk benefits to all the individuals in the mimicry ring and depends on the density of individuals sharing the same colouration, as well as on their harmfulness. As a facilitative interaction, Mullerian mimicry between sympatric species may participate in shaping community structure (Chazot et al., 2014; Elias et al., 2008), and in limiting local extinction risks (Boussens-Dumon & Llaurens, 2021). While mimicry was well-studied in neotropical butterflies (Bates, 1862), it remains surprisingly poorly studied in wasps and bees (Hymenoptera: Aculeata) (Willadsen, 2022), although these species are well-known for both their conspicuous coloration and their painful stings (Wallace, 1878). A few cases of convergent evolution of colour patterns are documented in bumblebees (*e*.*g*., Plowright & Owen, 1980; Williams, 2007) and velvet ants (*e*.*g*., Wilson et al., 2012, 2015), but these cases represent only a fraction of the wide variety of conspicuous patterns and painfully stinging found throughout the Aculeata clade. Mullerian mimicry within and among bees and wasps is indeed probably widespread. While most models of Mullerian mimicry consider equal levels of defence between the sexes, as observed in Lepidoptera, only females are defended in Aculeata. This female-limited defence may play a substantial role on population dynamics and may modulate the effect of Mullerian mimicry on community assembly and extinction risks in Aculeata. Here, we thus develop a mathematical model to investigate the effects of warning coloration and Mullerian mimicry on species co-existence in species where defences are restricted to a single sex, as observed in Aculeata.

In most Aculeata, females escape predators because of the pain induced by their sting and by the injected venom (Schmidt, 2004). The stinger in wasp and bee females may then induce a substantial difference in survival between defended females and undefended males, in habitats where predators are common. The relative abundance of defended females and undefended males sharing the same warning colours, whatever the species they belong to, then modulates the predation risk: the proportion of attacks tends to increase when the proportion of defended individuals decreases (Brower, 1960; Jones et al., 2013). However, this increase crucially depends on the levels of the noxiousness (Brower, 1960; Davis Rabosky et al., 2016; Howarth et al., 2004). Because of the lack of defences in males, the sex-ratio within populations of Aculeata is expected to be a key factor in shaping the individual predation risk within mimetic communities.

The sex-ratio of bees and wasps (haplodiploid species) is linked to the fertilization of the eggs, males being produced from an unfertilized egg. The sex-ratio of the offspring can be modulated by fertilized females, storing sperm cells after mating and thus regulating the proportion of fertilized eggs in their progeny. Males tend to be smaller than females, because size is a less important factor for male fitness (Stubblefield & Seger, 1994). Hence, assuming an equal investment of energy in each sex, we could observe biased sex-ratio in favour of the less expensive sex, namely the males (Trivers & Hare, 1976). Although, other factors may influence the sex-ratio in the progeny like seasonality, resource quality and quantity and population structure (Werren, 1987). Thus, the sex-ratio in the progeny produced by a female may vary between different species, depending on the relative investment in son production. Such variation can have a deep influence on adult sex-ratio in natural populations. Because of the sex-linked differences in defence, variations in sex-ratio in mimetic species may have a deep impact on individual survival, as well as on species extinction risk within mimicry rings.

Furthermore, in Aculeata, male and female can either display the same colour pattern or look very different, leading to important difference in individual predation risk and population dynamics. In species where males exhibit the same conspicuous pattern as females, they benefit from protection against predators due to mimicry towards the female signal. In contrast, striking sexual dimorphism in warning signals can be observed in other species (*e*.*g*., *Dasymutilla gloriosa*, Mutillidae; *Aplochares imitator*, Pompilidae) (Pitts & Sadler, 2017; Wilson et al., 2015). In these sexually-dimorphic species, males can display warning colours exhibited by females from other defended species living in sympatry (Evans, 1968), resulting in Batesian mimicry towards the defended species (Bates, 1862). This Dual Sex-limited Mimicry (DSLM) may have a contrasted effect on community assemblages. The effect of sex-ratio, as well as of the sexual dimorphism on population dynamics of mimetic species thus needs to be investigated to study the impact of these mutualistic interactions on extinction risk in Aculeata.

Using a differential equations model, we thus explicitly modelled the population dynamics of male and female populations of Aculeata assuming shared predator community and competition for resources. First, using a single species model, we explored the effect of sex-ratio and female noxiousness on local extinction risk. Then we built a two-species model to investigate the effect of mimicry on species persistence and co-existence, by specifically focusing on the effect of variations in female noxiousness and sex-ratio in the two interacting species. Finally, we explored the interaction between mimicry and sex-ratio in the species co-existence when dual sex-limited mimicry occurs between sympatric species.

## Material & Methods

To investigate the effect of mimicry on communities of sex-limited defended species, such as the Aculeata, we built a deterministic model considering population dynamics of both male and female of a haplodiploid species. To explore the effect of automimicry on extinction risk, we first studied a single species model. Then, we used a two-species model to test for the effect of mimicry between species in either both sexes or in males only (with a case of dual sex-limited mimicry). All variables and parameters used in these models are detailed in Table 1.

**Table 1.**
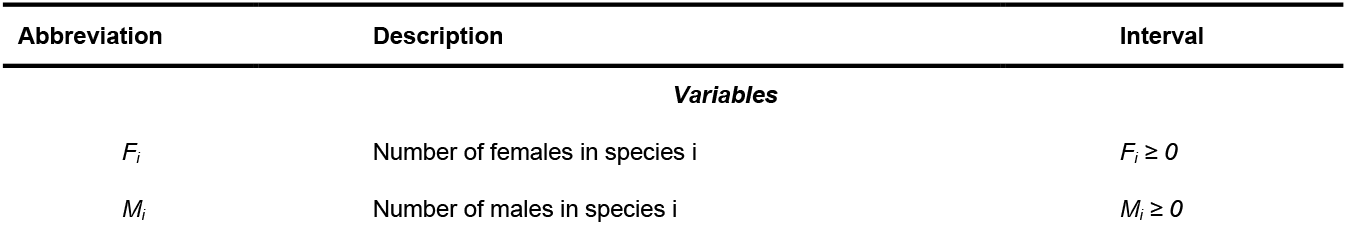
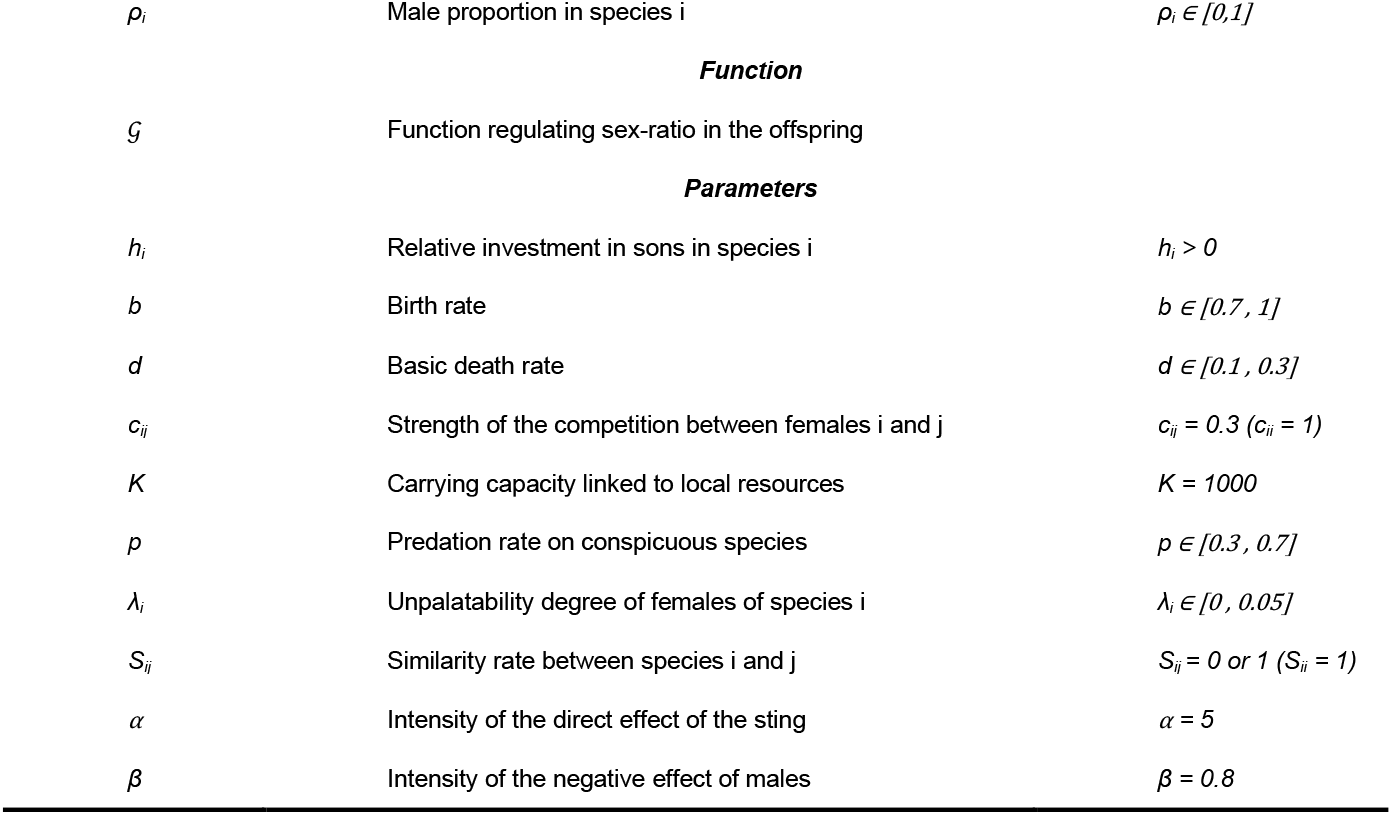
Variable and parameters, with their signification and values.

| Abbreviation | Description | Interval |
| --- | --- | --- |
| **Variables** |  |  |
| $F_i$ | Number of females in species $i$ | $F_i \geq 0$ |
| $M_i$ | Number of males in species $i$ | $M_i \geq 0$ |
| $\rho_i$ | Male proportion in species i | $\rho_i \in [0,1]$ |
| **Function** |  |  |
| $\mathcal{G}$ | Function regulating sex-ratio in the offspring | |
| **Parameters** |  |  |
| $h_i$ | Relative investment in sons in species i | $h_i > 0$ |
| $b$ | Birth rate | $b \in [0.7, 1]$ |
| $d$ | Basic death rate | $d \in [0.1, 0.3]$ |
| $c_{ij}$ | Strength of the competition between females i and j | $c_{ij} = 0.3$ ($c_{ii} = 1$) |
| $K$ | Carrying capacity linked to local resources | $K = 1000$ |
| $p$ | Predation rate on conspicuous species | $p \in [0.3, 0.7]$ |
| $\lambda_i$ | Unpalatability degree of females of species i | $\lambda_i \in [0, 0.05]$ |
| $S_{ij}$ | Similarity rate between species i and j | $S_{ij} = 0$ or $1$ ($S_{ii} = 1$) |
| $\alpha$ | Intensity of the direct effect of the sting | $\alpha = 5$ |
| $\beta$ | Intensity of the negative effect of males | $\beta = 0.8$ |

### 1. Model and assumptions

Let *F*_*i*_ and *M*_*i*_ be the population density of females and males from the species i respectively. The changes in male and female densities over time, noted *dF*_*i*_*/dt* and *dM*_*i*_*/dt* respectively, depend on the production of offspring of each sex (*O*_*i*_^*♀*^ and *O*_*i*_^*♂*^), competition between females (*C*_*i*_^*♀*^) and adult death. Adult death is composed of a basic mortality rate (*D*_*i*_^*♀*^ and *D*_*i*_^*♂*^) and a specific mortality rate caused from predation (*P*_*i*_^*♀*^ and *P*_*i*_^*♂*^). We thus denote:

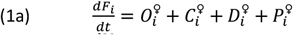

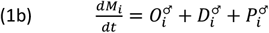

with *i* ∈ {1,2}.

#### 1.1. Offspring production

In order to define the sex-ratio at birth in the progeny of females, we used an increasing function bounded between 0 and 1, named *G (*based on Banks et al., 2017). This function determines the proportion of daughters in the offspring produced by females, depending on male proportion in the population. When the proportion of males increases in the population, the relative abundance of fertilised eggs (*i*.*e*., the proportion of daughters in the progeny) increases too. The intensity of this relationship is modulated by a parameter *h*_*i*_, which modulates the investment in son versus daughter production. When *h*_*i*_ *= 0*, sons are infinitely less costly than daughters and females produce only sons. Conversely, when *h*_*i*_ is high (*h*_*i*_ > 10), females produce only daughters. We chose values of *h*_*i*_ between 1 and 5 in order to explore sex-ratio from male-biased to female-biased. When the value of *h*_*i*_ increases, the quantity of fertilised eggs, given the proportion of males in the population noted *ρ*_*i*_, increases too:

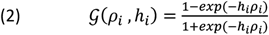

where *ρ*_*i*_ represents the proportion of males in the population and *h*_*i*_ the relative cost of producing sons in the species *i*.

The variation of population density (both female and males) due to offspring production by females is:

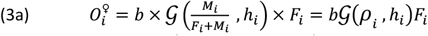

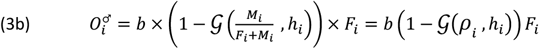

where *b* represents the individual rate at which females reproduce.

#### 1.2. Competition within and between species

Following existing models on population dynamics of mimetic species (*e*.*g*., Kumazawa et al., 2006; Sekimura et al., 2014; Yamauchi, 1994), we included exploitative competition in our model. We modelled competition only between females because most limiting resources of food and nesting sites are sought out only by females (Cane et al., 2017; Schneider et al., 2004). The effect of competition between females depends on two parameters: a coefficient of niche overlap *c*_*ij*_ between species *i* and *j*, and the limiting factor of resources *K* shared by sympatric species. When *j* = *i, c*_*ij*_ represents the strength of the intraspecific competition and we assumed *c*_*ii*_ = 1. Because we expected niche overlap to be maximum within species, interspecific competition is expected to be weaker than intraspecific one, so *c*_*ij*_ ≤ *1*. Except when explicitly mentioned, we considered *c*_*ij*_ *= 0*.*3* and *K = 1000*. The variation of female population density due to interspecific and intraspecific competition for resources is then:

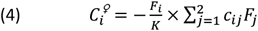

where *K* represents the carrying capacity linked to local resources and *c*_*ij*_ the coefficient of niche overlap between females *i* and *j*.

#### 1.3. Adult mortality

Males and females suffer from basic mortality (at rate *D*) and a mortality caused by predation (at rate *P*). The variation of female and male densities due to basic adult mortality are respectively:

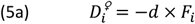

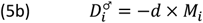

where *d* is the basic death rate.

Survival from predation then depend on the sex of the individual, because only female possess defences. The sting of female may facilitate their escape after an attack by a predator. We thus assumed that the mortality rate due to predation is different between males and females, considering females have a probability of escaping an attack depending on their noxiousness. Furthermore, survival from predation in both sexes can be increased because of predator learning. The predation terms for females and males can thus be written as:

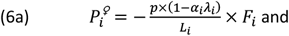

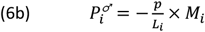

where *p* is the predation rate, *α*_*i*_ represents the direct effect of the sting on the probability for females to escape an attack, *λ*_i_ is the noxiousness of females and *L*_*i*_ represents the indirect protection due to mimicry. When *α*_*i*_ *= 0*, sting does not enhance female escaping, so that males and females have the same mortality rate due to predation.

Following Joron & Iwasa (2005) and suggested by Mallet & Joron (1999), we used a density-dependent effect of mimicry on predation. When a predator meets an unpalatable individual, it associates the noxiousness and the conspicuous pattern, reinforcing the protection provided by mimicry. As the number of unpalatable individuals sharing the same signal increases, the predation rate will decrease. Then, the predation death rate hyperbolically decreases as population size of defended prey increases. Note that this advantage against predators applies to both defended and undefended individuals (*i*.*e*., in both males and females in Aculeata), as long as they share the same conspicuous coloration. Nevertheless, the relative abundance of undefended and harmful individuals sharing the same conspicuous signal, respectively males and females in Aculeata, is likely to modulate the protection brought by mimicry: the proportion of attack tends to increase when the proportion of defended individuals decreases within a mimicry ring (Brower, 1960; Jones et al., 2013). Thus, we assumed that the proportion of males in a mimicry ring had a negative effect on protection provided by mimicry, so the indirect protection due to mimicry would be:

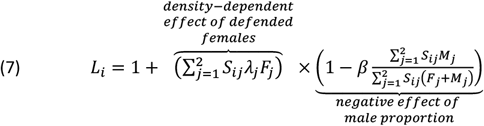

where *λ*_j_ is the noxiousness of female *j* and *S*_*ij*_ is the similarity rate of warning signals between species *i* and *j*. When *S*_*ij*_ *= 0*, there is no mimicry between individuals of species *i* and *j* while when *S*_*ij*_ *= 1*, the two species are perfect mimics (we leave aside cases of imperfect mimicry). Then *β* is the negative impact of harmless males on predator avoidance. When *β = 0*, there is no impact of undefended males on predator learning.

Finally, the variation of population density (both female and males) due to mortality caused by predation is:

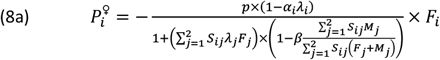

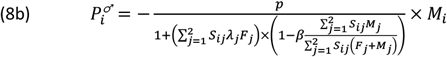

By combining equations (1a), (3a), (4), (5a), (8a) and (1b), (3b), (5b), (8b) we obtain the following system of two equations:

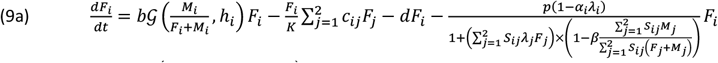

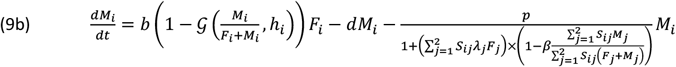

### 2. Numerical simulations

Except when explicitly mentioned, we randomly chose initial abundances (*F*_*i*_ and *M*_*i*_, which fix the initial male proportion *ρ*_*i*_), birth rate (b), death rate (d) and predation rate (p) in each simulation, and the other parameters were fixed to their default values (see Table 1). Very few ecological data are available in the literature to accurately estimate the values of most parameters, and some parameters might be difficult to directly measure in the wild (*e*.*g*., *λ, α* and *β*). Hence, the intervals explored and the fixed values were chosen based on previous exploratory simulations: we focused on parameters values enabling a large range of possible outcomes (*i*.*e*., values below or above these ranges force the maintenance or extinction of populations) to explore a diversity of ecological scenarios. Note that the absolute values considered might depend on the relationship between the parameters and the number of species studied.

#### 2.1. Exploring the effect of noxiousness and sex-ratio on extinction risks for one species

In mimetic populations, the protection against predation is based on the unpalatability of defended individuals and their relative abundance in the population. As a first step, we studied how these two aspects influence the defence level of a mimetic population as well as their extinction risks, considering only one species. From the general equations (9a) and (9b) we can write the change of female and male densities in a single species by fixing *F*_*2*_ *= 0* and *M*_*2*_ *= 0*. Thus, we obtain:

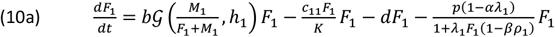

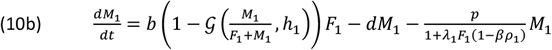

First, we explored the state of the population at equilibrium depending on predation pressure (*p*) on the one hand, and the two main components of the group defence, *i*.*e*., female noxiousness (*λ*) and sex-ratio (*i*.*e*., the proportion of defended females *vs*. harmless males, driven by the investment in sons *h*). Altough we only considered a single species, the system was difficult to study analytically because of the shape of the *G* function and the different effects of competition and predation on males and females (see the mathematical detail in supplementary). We therefore explored the single-species model numerically only by performing simulations for different values of *p* within [0,1] with a step of 0.1, and different values of *λ*_*1*_ within [0,0.05] with a step of 0.005. We recorded the state of the population at equilibrium (extinct or maintained) as well as the proportion of males, for 500 simulations per combinations of *p* and *λ*_*1*_, and this for two values of investment in sons: in favour of males (*h*_*1*_ *= 2*) or in favour of females (*h*_*1*_ *= 5*).

In addition, we also observed if mortality induced by predation has an effect on sex-ratio at equilibrium or if it remained constant. Thus, we performed 5000 simulations with random values of p, for 4 degrees of investment in sons (*h*_*1*_ ∈ *{2,3,4,5}* with a fixed value of *λ*_*1*_ *= 0*.*01*) and we recorded the male proportion at equilibrium. We made linear regressions and we tested the effect of predation pressure on the proportion of males at the equilibrium, using python packages *scikit-learn ver. 0*.*24*.*1* (Pedregosa et al., 2011) and *seaborn ver. 0*.*11*.*1* (Waskom et al., 2017).

Furthermore, we explored the effect of the direct protection provided by the sting for females (driven by *α*) as well as the cost of male proportion on predator learning (driven by *β*) on the population equilibrium. These two parameters are linked to the harmfulness of females and the investment in son production, and are specific to the Aculeata model. We chose values of *λ*_*1*_ *= 0*.*02* and *p = 0*.*6* for which the population was maintained in the first experiment, then we varied *α* within [0,10] with a step of 1, and *β* within [0,1] with a step of 0.1, for random values of *h*_*1*_ within [2,5]. We recorded the frequency of persistence at the equilibrium for 500 simulations per combinations of *α* and *β*.

#### 2.2. Investigating the effect of mimicry between two species

We investigated the effect of mimicry on co-existence of species in sympatry considering two species, mimetic or not (see the detailed systems of equations S1 and S2 in supplementary). To focus on the effect of mimicry, we considered females from both species as equally noxious (*λ*_*1*_ = *λ*_*2*_), and similar investment in male production in both species (*h*_*1*_ *= h*_*2*_). We performed different simulations with different values of *λ*_*1*_ *= λ*_*2*_ within [0.01, 0.05] using an increment of 0.005, and *h*_*1*_ *= h*_*2*_ within [1, 5] with a step of 0.5. We compared two types of community: either the two species display a different warning signal (no mimicry, *S*_***ij***_ = 0) or both species display the same warning signal (mimicry, *S*_*ij*_ = 1). We ran 500 simulations for each set of parameters and each type of community and we recorded the equilibrium state for each species. We then calculated the frequency of co-existence observed over the 500 simulations, for each combination of *λ* and *h* values.

#### 2.3. Investigating the level of mutualism between mimetic species on their co-existence

Because females of mimetic species contribute to the protection against predators, we tested the impact of uneven mutualistic interaction on species extinction and co-existence using unequal defence level between the species (*λ*_*1*_ ≠ *λ*_*2*_) and different investment in male production (*h*_*1*_ ≠ *h*_*2*_).

First, we explored uneven female noxiousness and investment in sons separately. We performed simulations with different values of *λ*_*1*_ and *λ*_*2*_ within [0, 0.05] with a step of 0.005, with random values of h_1_ = h_2_. In the same way, we performed simulations where the values of *h*_*1*_ and *h*_*2*_ varied within [1, 5] with a step of 0.5, with random values of *λ*_*1*_ *= λ*_*2*_. In either case, we recorded the equilibrium obtained from 500 simulations per combinations of *λ*_*1*_ and *λ*_*2*_ (or *h*_*1*_ and *h*_*2*_ respectively), for each community.

Then, we considered unequal female noxiousness (*λ*_*1*_ ≠ *λ*_*2*_) and different investment in male production (*h*_*1*_ ≠ *h*_*2*_) at the same time. We performed simulations with different values of *λ*_*1*_ and *λ*_*2*_ within [0.01, 0.05] with a step of 0.01, and of *h*_*1*_ and *h*_*2*_ within [1, 5] with a step of 1. We ran 500 simulations for each parameter set (*i*.*e*., combinations of *λ*_*1*_, *λ*_*2*_, *h*_*1*_ and *h*_*2*_ values) and recorded the equilibria for the two types of communities (either mimetic or not). In both experiments, we considered the equilibrium state at the scale of the community: either co-extinction, extinction of one species (1 or 2) or co-existence.

#### 2.4. Investigating the effect of dual sex-limited mimicry

Finally, we investigated the effect of dual sex-limited mimicry, considering that species 2 display sexual dimorphism in coloration, with males being mimetic to species 1. In contrast, the species 1 stayed monomorphic. We thus considered a slightly different model for indirect mimetic protection, by assigning different similarity rates *S*_*ij*_ for males and females:

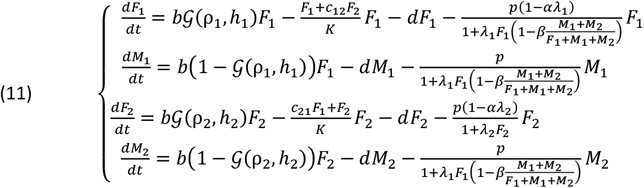

where 1 + *λ*_2_*F*_2_ is the indirect mimetic protection for females *F*_2_ and 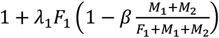 is the indirect mimetic protection for populations *F*_*1*_, *M*_*1*_ and *M*_*2*_.

With this model, we performed different simulations with different values of *λ*_*1*_ and *λ*_*2*_ within [0.01, 0.05] with a step of 0.01, and of *h*_*1*_ and *h*_*2*_ within [1, 5] with a step of 1. We ran 500 simulations for each parameter set (*i*.*e*., combinations of *λ*_*1*_, *λ*_*2*_, *h*_*1*_ and *h*_*2*_ values) and recorded the equilibrium for the community. Because female population of the species 1 has to carry the cost of the two male populations, we reduced the intensity of the cost of males on predator learning by fixing *β* = 0.5, instead of *β* = 0.8.

### 3. Running simulations

Simulations were performed using Python *ver. 3*.*8*.*8* (Van Rossum & Drake, 2009) and differential equations were solved using the function *odeint* from the package *Scipy ver. 1*.*6*.*2* (Virtanen et al., 2020). The scripts are available on the following link: https://zenodo.org/badge/latestdoi/553618533.

We ran simulations during a number *n* of time intervals with Δt = 50 for each interval and a time-step of 0.1, which makes 500 times values per interval. Simulations were stopped when population densities varied less than 10^−4^ between two time intervals. Then a population was considered extinct at equilibrium when female (and male) densities were under 10^−3^. Note that the equilibria obtained were the same when using the value 10^−6^ as a threshold.

The data were analysed and visualised using the packages *Pandas ver. 1.2.4* (McKinney et al., 2010) and *Matplotlib ver. 3*.*5*.*2* (Hunter, 2007).

## Results

### 1. Effect of female noxiousness and sex-ratio on the extinction risk for a single species

Mortality from predation depends on predation pressure (*p*) and the defence level at the scale of the mimetic population, which relies mainly on the proportion of females (driven by investment in sons *h*) and their noxiousness (*λ*). First, we studied the influence of these two components on the extinction risk for a single species.

Our simulations suggest that when the predation rate is high and the noxiousness of females is low, the species goes extinct (Figure 1). When the cost of producing sons is low with respect to daughters (*h*_*1*_ *= 2*, Figure 1a), the sex-ratio at equilibrium is male-biased. Extinction then occurs for lower values of predation, because the low density of females limits the protection against predators. Conversely, when producing sons is more costly (*h*_*1*_ *= 5*, Figure 1b), this favours the persistence of the species, even for high predation rate or limited female noxiousness (Figure 1b). Thus, a species producing a male-biased sex-ratio at birth could be more sensitive to extinction by predation.

**Figure 1.**
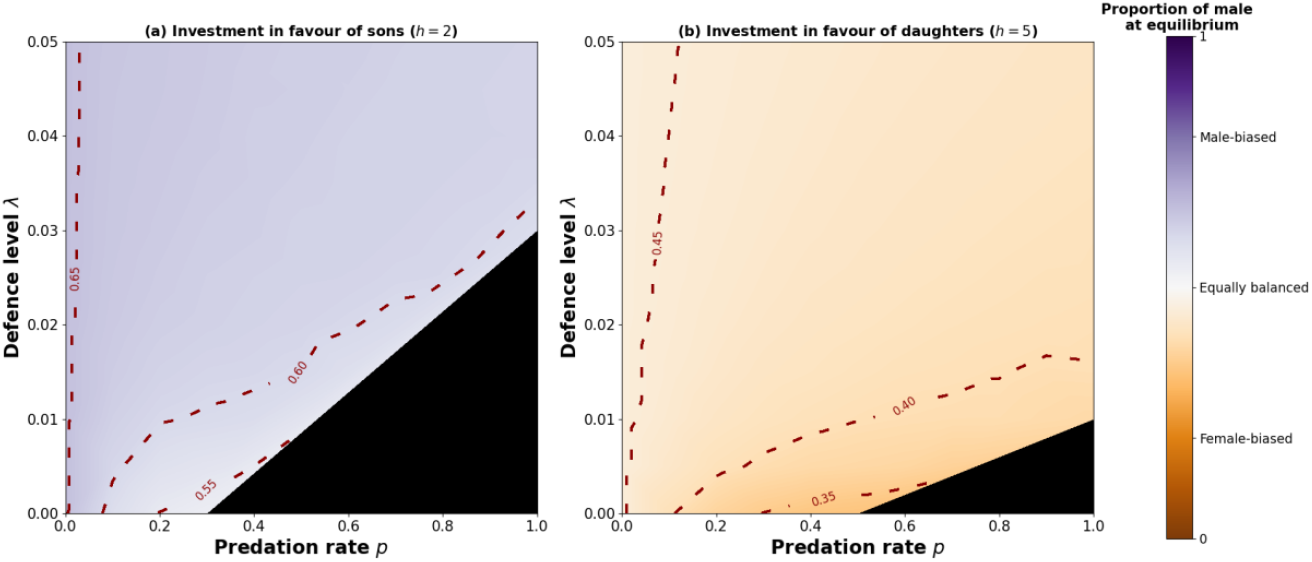
Effect of predation rate and female noxiousness on the persistence of the population at equilibrium, for two values of relative investment in sons: in favour of males (*h*_*1*_ *= 2*, figure 1a) or in favour of females (*h*_*1*_ *= 5*, figure 1b). The population is considered extinct when the equilibrium density is below 0.001 (black areas). In case of persistent population, the proportion of males at equilibrium averaged over 500 simulations is also represented: purple and orange colours indicate male and female-biased sex-ratio respectively. For each simulation, initial abundance, male proportion, birth rate, and death rate are chosen randomly and the other parameters are fixed (see Table 1).

Female noxiousness and sex-ratio both affect the defence level of the group and therefore the persistence of the mimetic population. The extinction risk is reduced when females are sufficiently noxious and abundant. The population can even be maintained if females are less numerous than males (male-biased sex-ratio), as long as they are sufficiently harmful (Figure 1a).

In addition to these two components, the presence of a stinger in females only also has an impact on extinction risk. The frequency of persistence of the population at equilibrium is lower when the cost of undefended males increases (*β* > 0 – Supplementary S3), but is higher when the survival advantage for females increases (*α* > 0 – Supplementary S3). Thus, the group defence level in species with female-limited defences mostly relies on the noxiousness of the individuals and their abundance. Yet, the cost of undefended individuals (specific component of automimetic populations) and the female-limited survival advantage of the stinger (specific to Aculeata species) modulate this defence.

Finally, linear regressions show a significant effect of predation on the proportion of males at equilibrium (estimate for *p*: −0.20, *F*_*1*_^*7827*^ *= 2082*.*86, p-value < 2*.*2e-16* – Supplementary S4 and S5). The proportion of male is always lower with predation than without predation (*p = 0* – Supplementary S4) and the sex-ratio tends to be equally balanced, even female-biased, when the predation pressure increases. When mortality increases due to higher predation rate, competition within females decreases due to fewer individuals. The increase in mortality is partly compensated by the decrease in competition, but only for females. The impact of mortality is thus relatively lower for females than for males, resulting in a diminution of male proportion.

### 2. Positive effect of mimicry on species co-existence

We explored the effect of mimicry between two species on their co-existence, according to their female noxiousness *λ*_*i*_ and relative investment in sons *h*_*i*_ (which drive the sex-ratio). We considered equal noxiousness (*λ*_*1*_ = *λ*_*2*_) and investment in sons (*h*_*1*_ *= h*_*2*_), and we compared a community without mimicry (*S*_*ij*_ *= 0*) and a community with mimicry (*S*_*ij*_ *= 1*).

The frequency of co-existence increases when noxiousness of females and their proportion in the offspring increase. Similarly to the single species model (Figure 1), these two components improve the defence level of the mimetic group and persistence of populations, and thus promote co-existence. However, for a given combination of *λ*_*1*_ *= λ*_*2*_ and *h*_*1*_ *= h*_*2*_, the frequency of co-existence at equilibrium is higher in the community with mimicry than without mimicry. In the mimetic community, co-existence is the most frequent equilibrium (observed on more than 50 % of the simulations - red line, Figure 2b) for smaller values of *λ*_*1*_ *= λ*_*2*_ and *h*_*1*_ *= h*_*2*_ than in the community without mimicry (Figure 2a). When the two species are strongly male-biased (*i*.*e*., when females are more costly than males to produce: *h*_*i*_ = 1, Figure 2), or when females are poorly noxious it increases the frequency of co-extinction.

**Figure 2.**
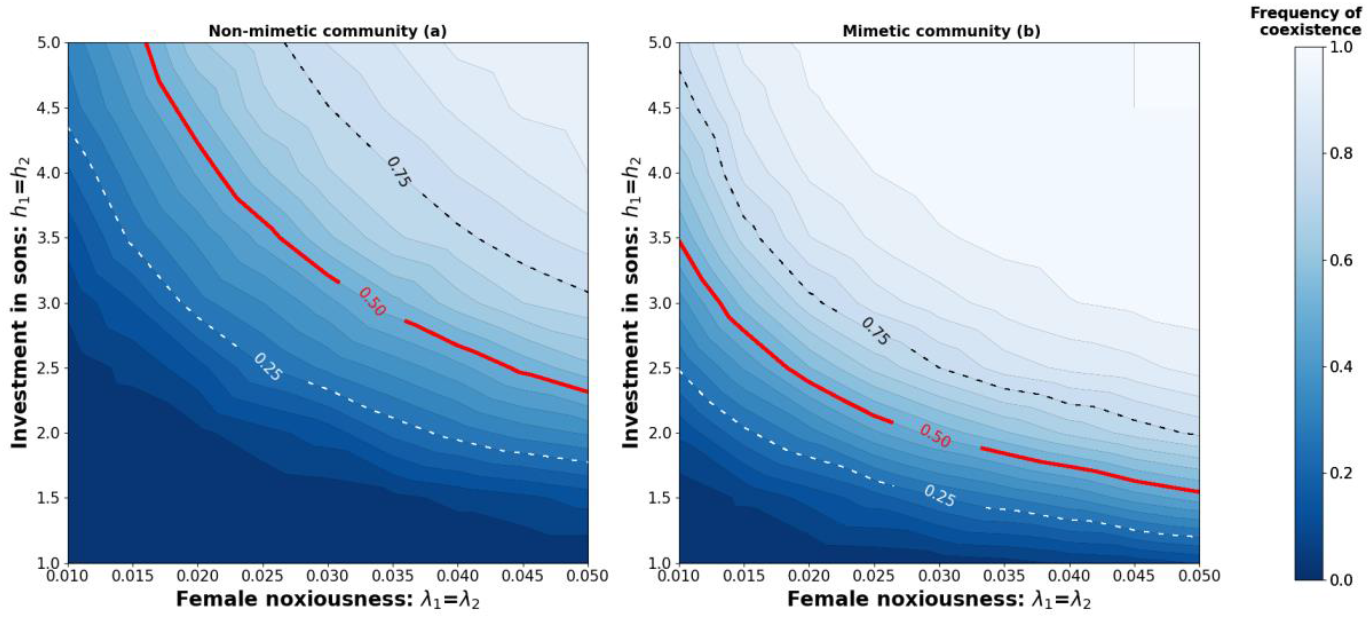
Effect of equal noxiousness and equal sex-ratio on the frequency of co-existence, for a community without mimicry (left side - Figure 2a) or with mimicry between species (right side - Figure 2b). The blue gradient represents the frequency of co-existence for 500 simulations. For each simulation, initial abundances, male proportions, birth rate, death rate and predation rate are chosen randomly and the other parameters are fixed (see Table 1). Lines represent equal levels of frequency: 50% for the red line, 25% for the light dotted line and 75% for the dark dotted line.

For a mimetic population or community to persist, it requires a minimum group defence level which mainly depends on female noxiousness and their abundance in the population. Considering equally harmful females in the two species, mimicry thus favours co-existence by increasing the abundance of defended individuals in the mimetic community.

### 3. Effect of uneven noxiousness and sex-ratio on the benefit of mimicry

Without mimicry, persistence of a species depends only on the noxiousness and relative abundance of their respective females. Co-existence is thus observed when both species populations have highly harmful females (high *λ*_i_ values - Figure 3b) and a low proportion of males (high *h*_*i*_ values - Figure 3c). When a species produces relatively more females and they are better defended than the other species, the most frequently observed equilibrium is the exclusion of the less protected species (purple and grey areas - Figure 3a). Co-extinction occurs when either a species has more females but poorly noxious, or the opposite (blue areas - Figure 3a).

**Figure 3.**
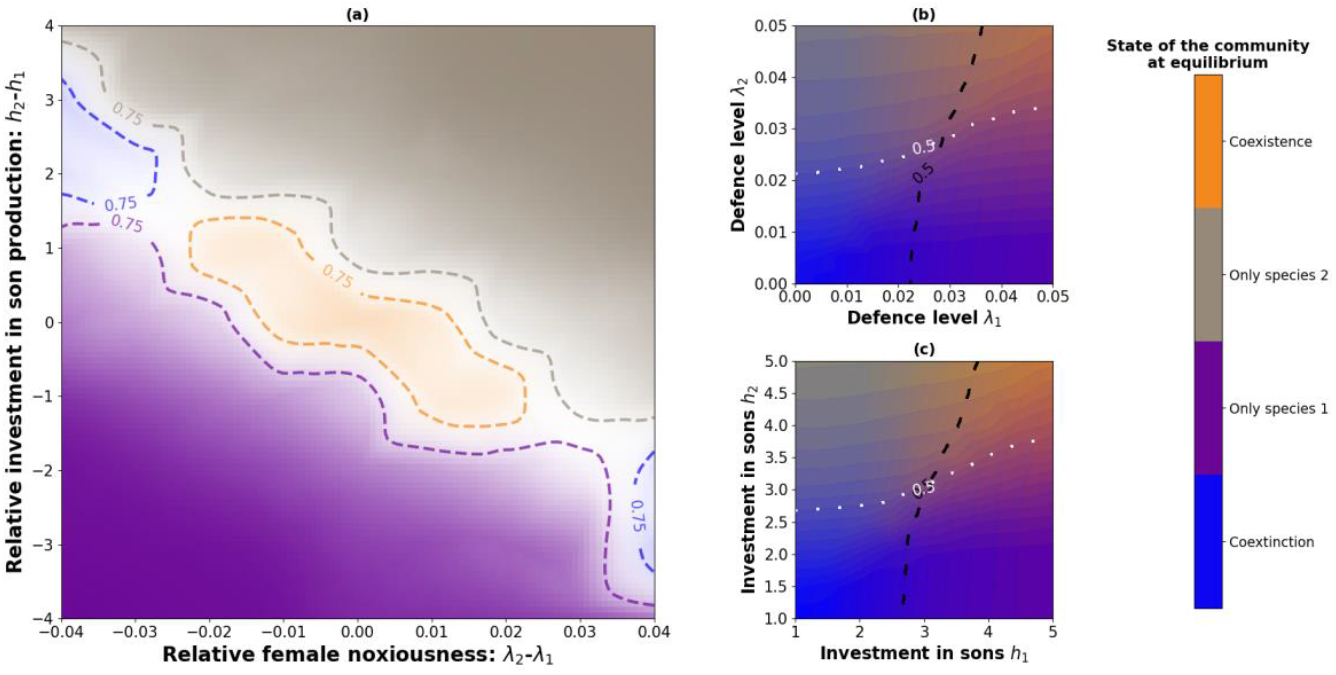
Effect of unequal female noxiousness and unequal investment in sons on the species co-existence for a non-mimetic community. We consider 4 equilibria: co-extinction (blue), co-existence (orange), only species 1 (purple) or only species 2 (grey). The colour gradient represents the frequency of equilibria for 500 simulations. In Figure 3a, because multiple pairs of parameters values may lead to the same value of *λ*_*2*_ *-λ*_*1*_ or *h*_*2*_ *-h*_*1*_, transparency levels match with the frequency of the most frequently observed equilibrium (full transparency corresponds to a frequency of 25% or under). For each simulation, initial abundances, male proportions, birth rate, death rate and predation rate are chosen randomly and the other parameters are fixed (see Table 1). Equal defence levels or sex-ratio are also randomly chosen when they are not plotted (for Figure 3b and 3c). The black and white lines represent the limit of 50% observed persistence, respectively for species 1 and 2.

When species are mimetic, the co-existence occurs as soon as a one species out of the two species is sufficiently protected (Figure 4a), either because their females are harmful (high *λ*_*i*_ values - Figure 4b) or relatively abundant (high *h*_*i*_ values promoting female-biased sex-ratio - Figure 4d). Harmless mimetic species can even be maintained (*λ*_*i*_ *= 0*), when the other species has very noxious females (Batesian mimicry). We see that co-existence occurs for most values of *λ*_*1*_, *λ*_*2*_, *h*_*1*_ *and h*_*2*_ (orange area, Figure 4). Species exclusion is still observed when the difference of investment in sons is important (*Δh = 4 or - 4* – Figure 4a) because the difference of female densities between the two species leads to competitive exclusion. Hence, mimicry favours co-existence in female-limited defence, even with unbalanced species traits

**Figure 4.**
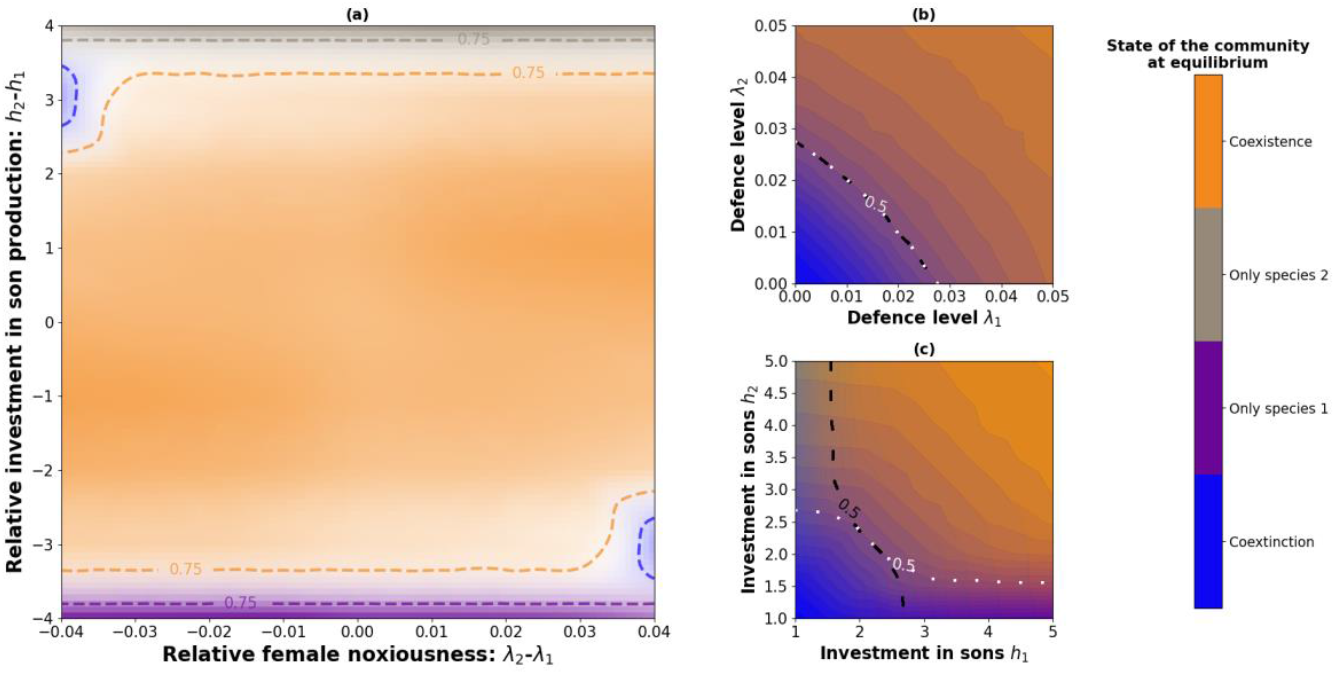
Effect of unequal female noxiousness and unequal investment in sons on the species co-existence for a mimetic community. We consider 4 equilibria: co-extinction (blue), co-existence (orange), only species 1 (purple) or only species 2 (grey). The colour gradient represents the frequency of equilibria for 500 simulations. In Figure 4a, because multiple pairs of parameters values may lead to the same value of *λ*_*2*_ *-λ*_*1*_ or *h*_*2*_ *-h*_*1*_, transparency levels match with the frequency of the most frequently observed equilibrium (full transparency corresponds to a frequency of 25% or under). For each simulation, initial abundances, male proportions, birth rate, death rate and predation rate are chosen randomly and the other parameters are fixed (see Table 1). Equal defence levels or sex-ratio are also randomly chosen when they are not plotted (for Figure 4b and 4c). The black and white lines represent the limit of 50% observed persistence, respectively for species 1 and 2.

### 4. Effect of dual sex-limited mimicry on co-existence

Finally, we explore the effect of dual sex-limited mimicry (DSLM) on species co-existence, considering the species 1 as monomorphic and the species 2 as dimorphic.

Our simulations show that species co-existence is frequent only in a restricted range of relative female noxiousness (*-0*.*04 < Δλ < 0*.*02* – Figure 5) and investment in sons (*-2*.*5 < Δh < 1*.*5* – Figure 5). These values correspond to situations where the monomorphic species is relatively better protected than the dimorphic species, either with more noxious and/or more abundant females (orange area – Figure 5).

**Figure 5.**
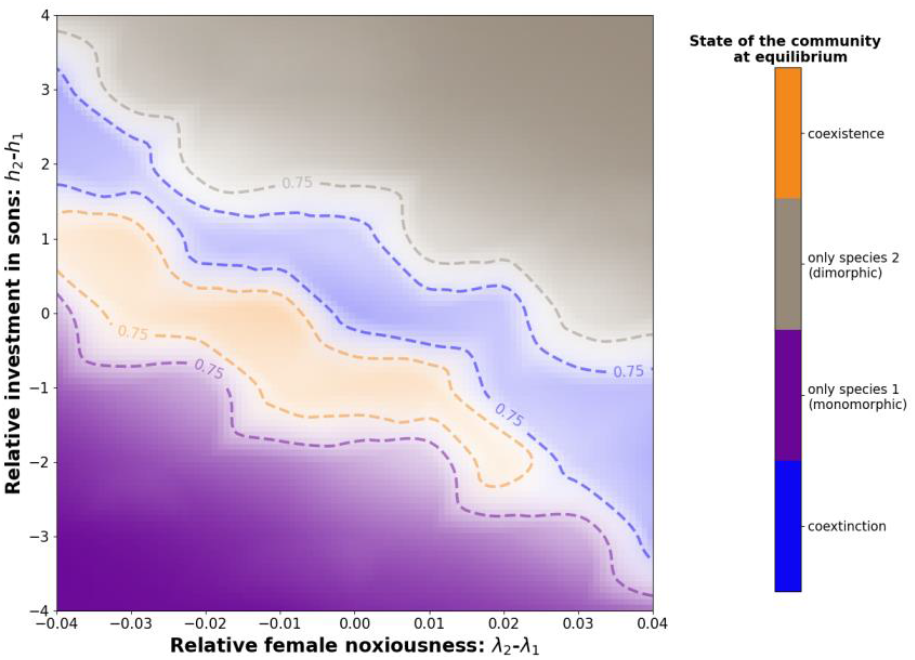
Effect of unequal female noxiousness and unequal investment in sons on the species co-existence, considering a monomorphic species (species 1) and a dimorphic species (species 2). Males of the second species mimic individuals of the species 1, while females are aposematic but with a distinct colour pattern. We consider 4 equilibria: co-extinction (blue), co-existence (orange), only species 1 (purple) and only species 2 (grey). Colour represents the most frequently observed equilibrium for 500 simulations. Because multiple pairs of parameters values may lead to the same value of *λ*_*2*_ *-λ*_*1*_ or *h*_*2*_ *-h*_*1*_, transparency levels match with the frequency of the equilibrium (full transparency corresponds to a frequency of 25% or under). For each simulation, initial abundances, male proportions, birth rate, death rate and predation rate are chosen randomly, the other parameters are fixed at their default value (see Table 1) except *β* = 0.5 in order to reduce the cost of males. Colored lines represent equal levels of frequency.

However, when both species have similar protections, the most frequent equilibrium is co-extinction. In these situations, females of the monomorphic species are not sufficiently protected from predation due to the cost of undefended males, leading to their extinction. Because males of the dimorphic species are no longer protected and cannot maintain themselves, the female population of this species will decrease until there are not enough defended individuals to ensure the protection of the colour pattern, leading to the extinction of the second species (blue area – Figure 5).

Considering dual sex-limited mimicry, females of the two species do not share the same aposematic pattern and therefore only interact negatively through competition. When a female population is better protected than the other one, this leads to species exclusion (purple and grey areas – Figure 5). The second species may persist without the other species protecting its males if dimorphic females are protected enough to survive despite the decrease of their population.

With a non-mimetic community, similar group defence levels between the two mimetic populations promote co-existence (Figure 3), but favour co-extinction when we consider a case of dual sex-limited mimicry (Figure 5). Instead, co-existence occurs when group defences levels are asymmetrical between the two populations and in favour of the monomorphic species, which carries the cost of all undefended males. Moreover, the co-existence is much less frequent in the community with dual sex-limited mimicry than with symmetric mimicry between the two species (Figure 4). Thus, dual sex-limited mimicry increases the risks of co-extinction, especially when both species have the same level of group defence. Under these conditions, co-existence requires a lower level of defence in the dimorphic species. In the absence of males mimicking females from species 2, limited abundance of species 2 reduces competition with species 1 females, and favour co-existence. Note that we reduced the cost of males on predator learning for these simulations (*β* = 0.5). With a value of *β* = 0.8, co-existence is frequent only when *Δλ = −0*.*01* and *Δh = −1*, so only when the monomorphic is slightly better protected than the other one.

## Discussion

In this paper, we provided a mathematical model for population dynamics of Mullerian mimetic species with female-limited defences, considering mimetic interaction between two species. Our findings are relevant to identify important ecological factors impacting the extinction risk in Aculeata communities.

### 1. Sex-ratio and extinction risk in Aculeata: the threat of male automimicry

Our model first considered the population dynamics in one haplodiploid aposematic species, where only females have defences and males act as Batesian mimics, specific to Aculeata. Our results showed that the resistance of such a species to an increasing predation pressure was related to two different components: the noxiousness of females, and the sex-ratio in the population. Our model highlighted the effect of the cost of investment in sons on the extinction risk in species with female-limited defence such as Aculeata species. Our results showed that for a fixed level of female noxiousness, the probability of extinction increases as the proportion of male increases in the population, when females are rarer than males (male-biased sex-ratio). Previous theoretical studies on Batesian mimicry complexes showed that the relative frequency of the mimics is correlated with the probability of a predator attack, when the model individuals are rarer than the mimics (Huheey, 1964; Holling, 1965; Emlen, 1968), and these results were supported by empirical studies (Lindström et al., 1997; Brower, 1960). However, the link between extinction risk and sex-ratio also depends on the level of unpalatability in females, which is consistent with the empirical (Lindström et al., 1997; Brower, 1960; Nonacs, 1985) and theoretical literature (Brower et al., 1970). Indeed, in Batesian mimicry, palatable mimics can be abundant when the level of noxiousness in the model species is high (Brower, 1960; Brower et al., 1970).

In solitary wasp and bee species, strongly male-biased, sex-ratio can be observed. Trivers & Hare (1976) indeed found male-biased sex-ratio for solitary wasps and bees from natural nests, bumblebees (*from* Webb, 1961 *in* Trivers & Hare, 1976) and some solitary species from trap nests (*from* Krombein, 1967 *in* Trivers & Hare, 1976), with sex-ratios with even more than two males per female in some species. Significant proportion of automimics have been reported by Brower (1969) in populations of the monarch butterfly *Danaus plexippus* (Lepidoptera), suggesting that important proportion of harmless individuals within population does not prevent the persistence of aposematic species in the wild.

The negative effect of males on the protection against predators can be reduced in species with sexually-differentiated phenology. In some Aculeata species, males come out after females during the season and therefore most predators have already learnt the aposematic signal. Waldbauer & Sheldon (1971) observed the phenology of Aculeata and of their insectivorous bird predators in a temperate area of the USA. The fledging of young birds mostly occurred during Summer and simultaneously with the abundance peak of Aculeata models, so the majority of naïve predator learning occurs during this period. Moreover, they also observed that stingless males were scarce in Aculeata populations during the summer and abundant in spring and fall. Longair (1981) and Seger (1983) both noted variations in the sex-ratio between the two generations of most bivoltine species of bees and wasps from temperate areas. The sex-ratio was balanced or female-biased for the summer generation, but becomes male-biased for the overwinter generation.

These empirical observations suggest that the lack of defence in aculeate males can influence population dynamics and may have influenced the evolution of investment in male offspring throughout the year. Thus, the extinction risk in Aculeata might depend on the variations of their sex-ratio through time in the different species, but also on their resemblance with other defended species living in sympatry.

### 2. Mimicry as a mutualistic interaction limiting extinction

Our results confirmed the positive effect of mimicry on species co-existence, despite the negative effects of undefended mimetic males and of the competition between females. Our model suggests that species co-existence depends on the level of noxiousness of females and on their proportion in the natural communities of mimetic species. Co-existence between two mimetic species may indeed occur when the level of defence of females from one species is sufficiently high, even if defences are lacking in the other species (*i*.*e*. Batesian mimicry).

Our results demonstrated the co-existence of mimetic species despite inter-specific competition. Co-mimetic species are found in sympatry, because the convergence evolution of warning coloration is promoted by the behaviour of the local predators feeding on these different aposematic species. Co-mimetic species therefore have largely overlapping ecological niches (Elias et al., 2008) and may thus often compete for resources. Interspecific competition tends to reduce species richness, but other ecological interactions have been documented to mediate the intensity of the competition. Models of foodweb indeed have shown that predation may reduce competition between prey (Droosel et al., 2001) and using a mathematical resource-consumer model, Gross (2008) has shown that positive interaction among exploitative competitors may enhance coexistence between species despite a net negative effect of interspecific interactions. For instance, co-existence in plant communities can be favoured through interactions that facilitate nutrient supply, either between plant species (Bertness & Leonard, 1997), *via* mycorrhizal interactions (Bergelson & Crawley, 1988) or through the effect of herbivores (Jensen & Nielsen, 1986). Our model highlights the mitigating effect of another mutualistic interaction, namely Mullerian mimicry, on the competitive exclusion between species arising from female competition for resources. Such a mitigating effect of Mullerian mimicry on species extinction risk was recently described in a previous model where equal level of defences were assumed across sexes (Boussens-Dumon & Llaurens, 2021). Our model demonstrates that, even when some mimetic individuals are unequally defended and therefore do not participate equitably in the predator education, Mullerian mimicry can still limit species exclusion caused by competition.

Our model considered the interaction between two species only, but natural communities of mimetic wasps and bees are composed of multiple species, occupy large geographical areas, and also interact with Batesian mimics. For instance, velvet ants and bumble bees are known to form large mimicry rings, in terms of number of species and geographical distributions (Hines et al., 2017; Wilson et al., 2015). Some conspicuous colour patterns are also widespread among Aculeata, and their persistence in large number of species might be promote by the positive effects of mimicry. The black-and-yellow pattern and the black-orange-black pattern are two common colourations among Aculeata and Hymenoptera in general (Boppré et al., 2016; Mora & Henson, 2019). Wasps and bees colour patterns occur also in other taxa of insects including undefended species like flies of the family Syrphidae (Leavey et al., 2021; Waldbauer, 1970). Thus, the protection provided by mimetic interaction involving Aculeata could benefit a large number of species and limit their extinction risk.

Mimicry between wasps and bees is a relevant factor to better understand the population dynamics and co-existence of Aculeata species. More broadly, since Aculeata are important pollinators, as are some of their Batesian mimics such as hoverflies (Syrphidae; Doyle et al., 2020), the positive effect of mimicry on co-existence could be even more important to consider given the current decline in pollinator populations (Biesmeijer et al., 2006; Hallman et al., 2017).

### 3. Male-limited mimicry as a destabilising factor in Aculeata communities

While our model generally suggests a positive effect of mimicry on species co-existence in Aculeata communities, the specific case of dual sex-limited mimicry (Evans, 1968) provides more contrasted result. In our model, the dual sex-limited mimicry (DSLM), where harmless males from a sexually dimorphic species resemble to defended females from another species, tends to increase the risks of co-extinction. Co-existence is indeed predicted in only a restricted range of female noxiousness and investment in sons: the monomorphic species mimicked by males from the other species needs to be relatively more protected than the dimorphic species, either with females more defended or more abundant, in order to maintain a sufficient level of protection, despite the cost of the additional mimetic males on the warning signal.

In Aculeata, DSLM was described in a few species of Pompilidae (Evans, 1968; Pitts & Sadler, 2017) and Mutillidae (Wilson et al., 2015). Other cases of DSLM may occur in Aculeata, especially for the mutilid wasps where the extreme sexual dimorphism probably prevent generalization of warning signals displayed by males and females (Pilgrim & Pitts, 2006). The evolution of colour dimorphism have been suggested to stem from behavioural differences between sexes (Heal, 1981; Van-Wright, 1971) and/or microhabitat divergence between male and female, resulting in contrasted selective pressures acting on either sexes. For instance, in the genus *Chirodamus* (Pompilidae), females hunt spiders on the ground like other wasp species including *Pepsis sp*., while males spend many times flying among social wasp workers (Ewans, 1968). In mutillid wasps, all females are apterous, while males do have wings and may have wider distribution areas and share the environment with other species. In wasps and bees, the obligatory sexual dimorphism in defences might also contribute to contrasted selection acting on male and female coloration and influence the evolution of dual sex-limited mimicry.

Our results highlight the impact of dual sex-limited mimicry on co-existence in Aculeata species. Undefended males are likely to represent a cost and might increase the extinction risk of the population, especially in species with poorly defended females or with a male-biased sex-ratio. In case of DSLM, species co-existence might stem from a precarious equilibrium so that anthropic pressures disturbing natural population dynamics of wasps and bees might have an even more significant effect on extinction risk than in other cases of mimicry between monomorphic species.

## Acknowledgements

The authors would like to thank Paul Chatelain for stimulating discussions on mimicry in Aculeata and for his help throughout the project, as well as the whole ‘Evolution and Development of Phenotypic Variations’ (ISYEB) for their feedback. We are also grateful to Thierry Spataro (iEES Paris) for his feedback on the first version of the model.

## Data, scripts, code, and supplementary information availability

Scripts and code are available online: https://zenodo.org/badge/latestdoi/553618533.

## Supplementary information

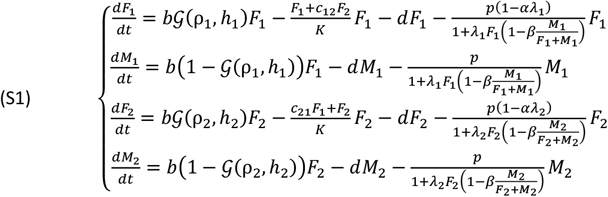

### S1. System of equations for the non-mimetic community

with *c*_*11*_ *= 1, c*_*22*_ *= 1, S*_*11*_ *= 1, S*_*12*_ *= 0, S*_*21*_ *= 0* and *S*_*22*_ *= 1*.

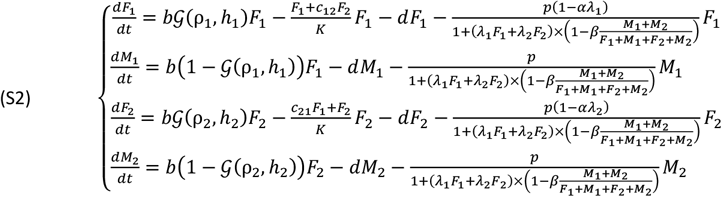

### S2. System of equations for the mimetic community

with *c*_*11*_ *= 1, c*_*22*_ *= 1, S*_*11*_ *= 1, S*_*12*_ *= 1, S*_*21*_ *= 1* and *S*_*22*_ *= 1*.

**Figure S3.**
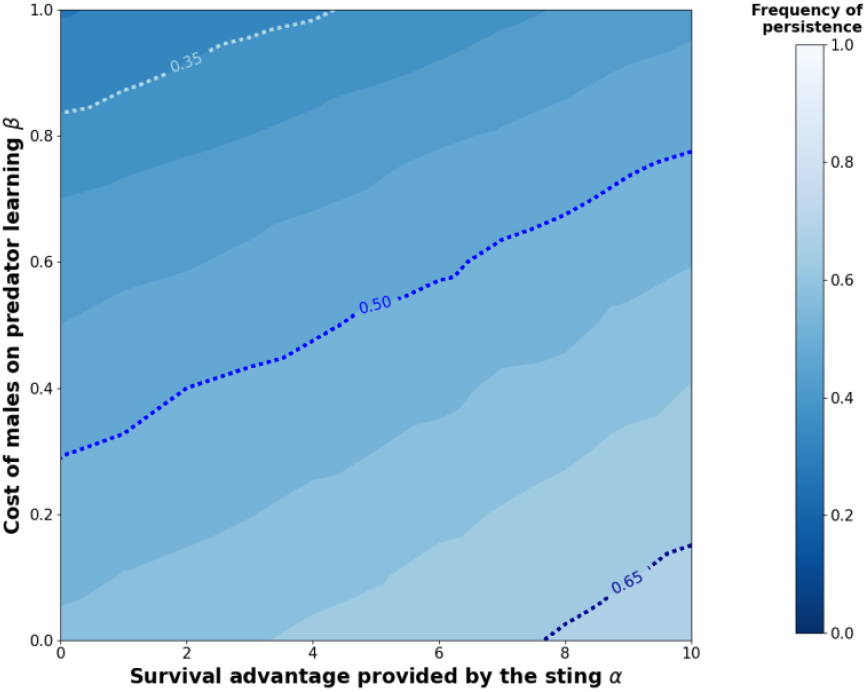
Effect of the female-limited survival advantage of the sting and of the cost of undefended males on the equilibrium. Simulations were run assuming *λ*_*1*_ *= 0*.*02* and *p = 0*.*6*. These parameter values insured the persistence of the population (see Figure 1). The frequency of persistence was averaged over 500 simulations, with random values of *h*_*1*_ within [2,5] for each simulation. Moreover, initial abundance, male proportion, birth rate, and death rate are chosen randomly and the other parameters are fixed (see Table 1). Blue dotted lines indicate equal levels of frequency (0.35, 0.5 and 0.65).

**Figure S4.**
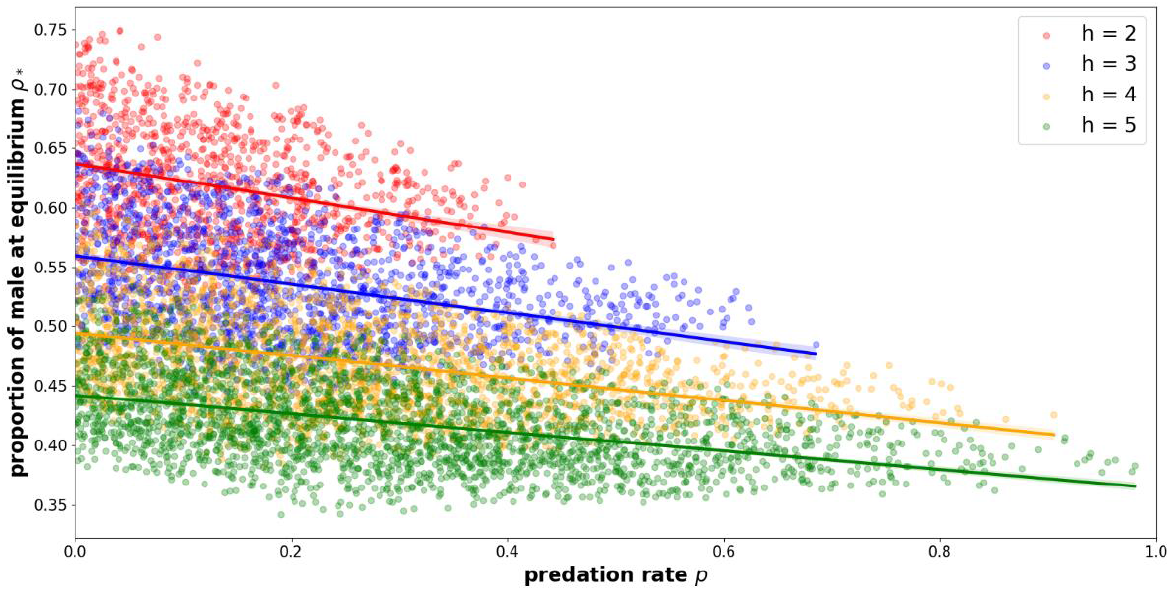
Effect of predation rate on male proportion at equilibrium, for different values of *h*. We made 5000 simulations for each value of *h*, with random values of p and a fixed value of *λ = 0*.*01*. Simulations leading to the extinction of the population are not represented. Moreover, initial abundance, male proportion, birth rate and death rate are chosen randomly and the other parameters are fixed (see Table 1).

**Figure S5.**
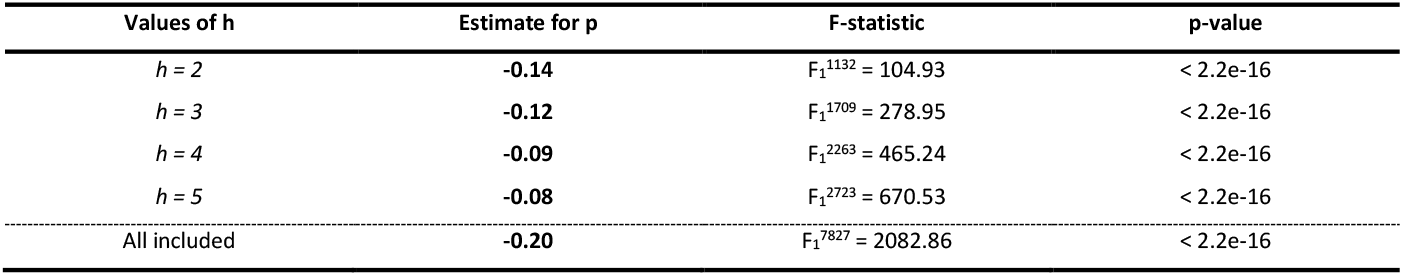
Effect of predation rate on male proportion at equilibrium for one species. We made 5000 simulations for each value of *h*, with random values of p and a fixed value of *λ = 0*.*01*. Simulations leading to the extinction of the population are not represented. Moreover, initial abundance, male proportion, birth rate and death rate are chosen randomly and the other parameters are fixed (see Table 1). Linear regressions were performed using python packages *scikit-learn ver. 0*.*24*.*1* (Pedregosa et al., 2011) and *seaborn ver. 0*.*11*.*1* (Waskom et al., 2017).

### S6. Study of the system (10a) and (10b)

We would like to stress that even if the dynamical system has only two coordinates, it is highly nonlinear and therefore difficult to study theoretically. The main difficulty derives from the function *G* appearing in birth rate, and the different effects on males and females of competition and predation.

Let us recall that the system describes the dynamics of *F*(*t*), *M*(*t*) the density of females and males and depends on the male ratio *ρ*(*t*) = *M*(*t*)/(F(*t*) + *M*(*t*)). The dynamical system writes

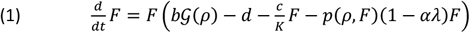

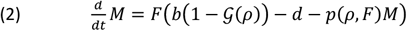

where 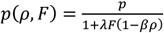.

We aim at characterizing the positive equilibria (F^*^, *M*^*^) of the system when it exists. Here we will actually compute *F*^*^ and *ρ*^*^ = *M*^*^/(F^*^ + *M*^*^), and we can then retrieve *M*^*^ = *ρ*^*^*F*^*^/(1 − *ρ*^*^).

By considering the total population size, we obtain that at equilibrium

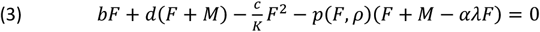

which leads dividing by (F + *M*)

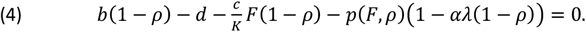

and thus

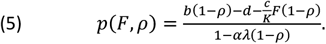

therefore

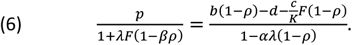

Using (5) in (1) we deduce that at equilibrium

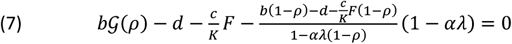

which reads

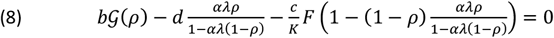

This allows to obtain *F*^*^ as a function of *ρ*^*^

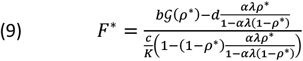

We can then replace *F*^*^ in (6) and obtain that *ρ*^*^ is a solution of

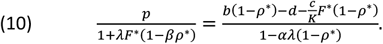

We see here, that due to the function *G* involved, and the non-linearity, an explicit expression for *ρ*^*^ is not available. Moreover, it is difficult to ensure that a solution *ρ*^*^ actually exists in (0,1) and that it gives a positive *F*^*^ in (9).

## References

Banks, H. T., Banks, J. E., Bommarco, R., Laubmeier, A. N., Myers, N. J., Rundlöf, M., & Tillman, K. (2017). Modeling bumble bee population dynamics with delay differential equations. Ecological Modelling, 351, 14–23. 10.1016/j.ecolmodel.2017.02.011

Bastolla, U., Fortuna, M. A., Pascual-García, A., Ferrera, A., Luque, B., & Bascompte, J. (2009). The architecture of mutualistic networks minimizes competition and increases biodiversity. Nature, 458(7241), 1018–1020. 10.1038/nature07950

Bates, H. W. (1862). XXXII. Contributions to an Insect Fauna of the Amazon Valley. Lepidoptera: Heliconidae. Transactions of the Linnean Society of London, 23(3), 495–566. 10.1111/j.1096-3642.1860.tb00146.x

Bergelson, J. M., & Crawley, M. J. (1988). Mycorrhizal infection and plant species diversity. Nature, 334(6179), 202–202. 10.1038/334202a0

Bertness, M. D., & Callaway, R. (1994). Positive interactions in communities. Trends in Ecology & Evolution, 9(5), 191–193. 10.1016/0169-5347(94)90088-4

Bertness, M. D., & Leonard, G. H. (1997). The Role of Positive Interactions in Communities: Lessons from Intertidal Habitats. Ecology, 78(7), 1976–1989. 10.1890/0012-9658(1997)078[1976:TROPII]2.0.CO;2

Biesmeijer, J. C., Roberts, S. P. M., Reemer, M., Ohlemüller, R., Edwards, M., Peeters, T., Schaffers, A. P., Potts, S. G., Kleukers, R., Thomas, C. D., Settele, J., & Kunin, W. E. (2006). Parallel Declines in Pollinators and Insect-Pollinated Plants in Britain and the Netherlands. Science, 313(5785), 351–354. 10.1126/science.1127863

Boppré, M., Vane-Wright, R. I., & Wickler, W. (2017). A hypothesis to explain accuracy of wasp resemblances. Ecology and Evolution, 7(1), 73–81. 10.1002/ece3.2586

Boussens-Dumon, G., & Llaurens, V. (2021). Sex, competition and mimicry: An eco-evolutionary model reveals unexpected impacts of ecological interactions on the evolution of phenotypes in sympatry. Oikos, 130(11), 2028–2039. 10.1111/oik.08139

Boutin, M. (2022). Maxime-Btn/Aculeata_Mimicry_Sexratio: v.1.2.0 (v.1.2.0). Zenodo. https://zenodo.org/badge/latestdoi/553618533

Bronstein, J. L. (1994). Our Current Understanding of Mutualism. The Quarterly Review of Biology, 69(1), 31–51. 10.1086/418432

Brower. (1960). Experimental Studies of Mimicry. IV. The Reactions of Starlings to Different Proportions of Models and Mimics. The American Naturalist, 94(877), 271–282. 10.1086/282128

Brower, L. P. (1969). ECOLOGICAL CHEMISTRY. Scientific American, 220(2), 22–29.

Brower, L. P., Pough, F. H., & Meck, H. R. (1970). THEORETICAL INVESTIGATIONS OF AUTOMIMICRY, I. SINGLE TRIAL LEARNING. Proceedings of the National Academy of Sciences, 66(4), 1059–1066. 10.1073/pnas.66.4.1059

Brower, L. P., van Brower, J., & Corvino, J. M. (1967). Plant poisons in a terrestrial food chain. Proceedings of the National Academy of Sciences, 57(4), 893–898. 10.1073/pnas.57.4.893

Bruno, J. F., Stachowicz, J. J., & Bertness, M. D. (2003). Inclusion of facilitation into ecological theory. Trends in Ecology & Evolution, 18(3), 119–125. 10.1016/S0169-5347(02)00045-9

Cane, J. H., Dobson, H. E. M., & Boyer, B. (2017). Timing and size of daily pollen meals eaten by adult females of a solitary bee (Nomia melanderi) (Apiformes: Halictidae). Apidologie, 48(1), 17–30. 10.1007/s13592-016-0444-8

Chazot, N., Willmott, K. R., Santacruz Endara, P. G., Toporov, A., Hill, R. I., Jiggins, C. D., & Elias, M. (2014). Mutualistic Mimicry and Filtering by Altitude Shape the Structure of Andean Butterfly Communities. The American Naturalist, 183(1), 26–39. 10.1086/674100

Chesson, P. (2000). Mechanisms of Maintenance of Species Diversity. Annual Review of Ecology and Systematics, 31(1), 343–366. 10.1146/annurev.ecolsys.31.1.343

Davis Rabosky, A. R., Cox, C. L., Rabosky, D. L., Title, P. O., Holmes, I. A., Feldman, A., & McGuire, J. A. (2016). Coral snakes predict the evolution of mimicry across New World snakes. Nature Communications, 7(1), 11484. 10.1038/ncomms11484

Doyle, T., Hawkes, W. L. S., Massy, R., Powney, G. D., Menz, M. H. M., & Wotton, K. R. (2020). Pollination by hoverflies in the Anthropocene. Proceedings of the Royal Society B: Biological Sciences, 287(1927), 20200508. 10.1098/rspb.2020.0508

Drossel, B., Higgs, P. G., & Mckane, A. J. (2001). The Influence of Predator–Prey Population Dynamics on the Long-term Evolution of Food Web Structure. Journal of Theoretical Biology, 208(1), 91–107. 10.1006/jtbi.2000.2203

Dunn, R. R., Harris, N. C., Colwell, R. K., Koh, L. P., & Sodhi, N. S. (2009). The sixth mass coextinction: Are most endangered species parasites and mutualists? Proceedings of the Royal Society B: Biological Sciences, 276(1670), 3037–3045. 10.1098/rspb.2009.0413

Edgar, J. A., Cockrum, P. A., & Frahn, J. L. (1976). Pyrrolizidine alkaloids inDanaus plexippus L. andDanaus chrysippus L. Experientia, 32(12), 1535–1537. 10.1007/BF01924437

Elias, M., Gompert, Z., Jiggins, C., & Willmott, K. (2008). Mutualistic Interactions Drive Ecological Niche Convergence in a Diverse Butterfly Community. PLoS Biology, 6(12), e300. 10.1371/journal.pbio.0060300

Emlen, J. M. (1968). Batesian Mimicry: A Preliminary Theoretical Investigation of Quantitative Aspects. The American Naturalist, 102(925), 235–241.

Evans, H. E. (1968). STUDIES ON NEOTROPICAL POMPILIDA (HYMENOPTERA) IV. EXAMPLES OF DUAL SEX-LIMITED MIMICRY IN CHIRODAMUS. 75(1), 1–22.

Gross, K. (2008). Positive interactions among competitors can produce species-rich communities. Ecology Letters, 11(9), 929–936. 10.1111/j.1461-0248.2008.01204.x

Hallmann, C. A., Sorg, M., Jongejans, E., Siepel, H., Hofland, N., Schwan, H., Stenmans, W., Müller, A., Sumser, H., Hörren, T., Goulson, D., & Kroon, H. de. (2017). More than 75 percent decline over 27 years in total flying insect biomass in protected areas. PLOS ONE, 12(10), e0185809. 10.1371/journal.pone.0185809

Hay. (1986). Associational Plant Defenses and the Maintenance of Species Diversity: Turning Competitors Into Accomplices. 26.

Heal, J. R. (1981). Colour patterns of Syrphidae. III. Sexual dimorphism in Eristalis arbustorum. Ecological Entomology, 6(2), 119–127. 10.1111/j.1365-2311.1981.tb00600.x

Hines, H. M., Witkowski, P., Wilson, J. S., & Wakamatsu, K. (2017). Melanic variation underlies aposematic color variation in two hymenopteran mimicry systems. PLOS ONE, 12(7), e0182135. 10.1371/journal.pone.0182135

Holling, C. S. (1965). The Functional Response of Predators to Prey Density and its Role in Mimicry and Population Regulation. The Memoirs of the Entomological Society of Canada, 97(S45), 5–60. 10.4039/entm9745fv

Holt, R. D. (2013). Species Coexistence. In S. A. Levin, Encyclopedia of Biodiversity (p. 667–678). Elsevier. 10.1016/B978-0-12-384719-5.00025-3

Howarth, B., Edmunds, M., & Gilbert, F. (2004). DOES THE ABUNDANCE OF HOVERFLY (SYRPHIDAE) MIMICS DEPEND ON THE NUMBERS OF THEIR HYMENOPTERAN MODELS? Evolution, 58(2), 367–375. 10.1111/j.0014-3820.2004.tb01652.x

Huheey, J. E. (1964). Studies of Warning Coloration and Mimicry. IV. A. Mathematical Model of Model-Mimic Frequencies. Ecology, 45(1), 185–188. 10.2307/1937125

Hunter, J. D. (2007). Matplotlib: A 2D Graphics Environment. Computing in Science & Engineering, 9(3), 90–95. 10.1109/MCSE.2007.55

Jensen, T. S., & Nielsen, O. F. (1986). Rodents as seed dispersers in a heath—Oak wood succession. Oecologia, 70(2), 214–221. 10.1007/BF00379242

Jones, R. S., Davis, S. C., & Speed, M. P. (2013). Defence Cheats Can Degrade Protection of Chemically Defended Prey. Ethology, 119(1), 52–57. 10.1111/eth.12036

Joron, M., & Iwasa, Y. (2005). The evolution of a Mullerian mimic in a spatially distributed community. Journal of Theoretical Biology, 237(1), 87–103. 10.1016/j.jtbi.2005.04.005

Kéfi, S., Baalen, M. van, Rietkerk, M., & Loreau, M. (2008). Evolution of Local Facilitation in Arid Ecosystems. The American Naturalist, 172(1), E1–E17. 10.1086/588066

Koh, L. P., Dunn, R. R., Sodhi, N. S., Colwell, R. K., Proctor, H. C., & Smith, V. S. (2004). Species Coextinctions and the Biodiversity Crisis. Science, 305(5690), 1632–1634. 10.1126/science.1101101

Kumazawa, F., Asami, T., Hayashi, T., & Yoshimura, J. (2006). Population dynamics of Batesian mimicry under interspecific competition. 14.

Kunte, K. (2009). The Diversity and Evolution of Batesian Mimicry in Papilio Swallowtail Butterflies. Evolution, 63(10), 2707–2716. 10.1111/j.1558-5646.2009.00752.x

Leavey, A., Taylor, C. H., Symonds, M. R. E., Gilbert, F., & Reader, T. (2021). Mapping the evolution of accurate Batesian mimicry of social wasps in hoverflies. Evolution, 75(11), 2802–2815. 10.1111/evo.14336

Lindström, L., Alatalo, R. V., & Mappes, J. (1997). Imperfect Batesian mimicry—The effects of the frequency and the distastefulness of the model. Proceedings of the Royal Society of London. Series B: Biological Sciences, 264(1379), 149–153. 10.1098/rspb.1997.0022

Longair, R. W. (1981). Sex Ratio Variations in Xylophilous Aculeate hymenoptera. Evolution, 35(3), 597–600. 10.2307/2408206

Mallet, J., & Joron, M. (1999). Evolution of Diversity in Warning Color and Mimicry: Polymorphisms, Shifting Balance, and Speciation. Annual Review of Ecology and Systematics, 30(1), 201–233. 10.1146/annurev.ecolsys.30.1.201

McKinney, W. (2010). Data Structures for Statistical Computing in Python. Proceedings of the 9th Python in Science Conference, 56-61. 10.25080/Majora-92bf1922-00a

Moeller, D. A. (2004). FACILITATIVE INTERACTIONS AMONG PLANTS VIA SHARED POLLINATORS. Ecology, 85(12), 3289–3301. 10.1890/03-0810

Mora, R., & Hanson, P. E. (2019). Widespread Occurrence of Black-Orange-Black Color Pattern in Hymenoptera. Journal of Insect Science, 19(2), 13. 10.1093/jisesa/iez021

Muller, F. (1879). Ituna and Thyridia: A remarkable case of mimicry in butterflies. Proc Entomol Soc Lond, xx–xxiv.

Nonacs, P. (1985). Foraging in a Dynamic Mimicry Complex. The American Naturalist, 126(2), 165–180. 10.1086/284407

Pedregosa, F., Varoquaux, G., Gramfort, A., Michel, V., Thirion, B., Grisel, O., Blondel, M., Prettenhofer, P., Weiss, R., Dubourg, V., Vanderplas, J., Passos, A., Cournapeau, D., Brucher, M., Perrot, M., & Duchesnay, É. (2011). Scikit-learn: Machine Learning in Python. Journal of Machine Learning Research, 12(85), 2825–2830.

Pilgrim, E. M., & Pitts, J. P. (2006). A Molecular Method for Associating the Dimorphic Sexes of Velvet Ants (Hymenoptera: Mutillidae). Journal of the Kansas Entomological Society, 79(3), 222–230. 10.2317/0511.09.1

Pitts, J. P., & Sadler, E. (2017). Association and description of the male of Aplochares imitator (Smith) (Hymenoptera: Pompilidae). Zootaxa, 4300(1), 135. 10.11646/zootaxa.4300.1.8

Plowright, R. C., & Owen, R. E. (1980). The Evolutionary Significance of Bumble Bee Color Patterns: A Mimetic Interpretation. Evolution, 34(4), 622. 10.2307/2408017

Raup, D. M. (1994). The role of extinction in evolution. Proceedings of the National Academy of Sciences, 91(15), 6758–6763. 10.1073/pnas.91.15.6758

Rowland, H. M., Ihalainen, E., Lindström, L., Mappes, J., & Speed, M. P. (2007). Co-mimics have a mutualistic relationship despite unequal defences. Nature, 448(7149), 64–67. 10.1038/nature05899

Rowland, H. M., Mappes, J., Ruxton, G. D., & Speed, M. P. (2010). Mimicry between unequally defended prey can be parasitic: Evidence for quasi-Batesian mimicry. Ecology Letters, 13(12), 1494–1502. 10.1111/j.1461-0248.2010.01539.x

Schmidt, J. O. (2004). Venom and the Good Life in Tarantula Hawks (Hymenoptera: Pompilidae): How to Eat, Not Be Eaten, and Live Long. Journal of the Kansas Entomological Society, 77(4), 402–413.

Schneider, S. S., Deeby, T., Gilley, D. C., & DeGrandi-Hoffman, G. (2004). Seasonal nest usurpation of European colonies by African swarms in Arizona, USA. Insectes Sociaux, 51(4), 359–364. 10.1007/s00040-004-0753-1

Seger, J. (1983). Partial bivoltinism may cause alternating sex-ratio biases that favour eusociality. Nature, 301(5895), 59–62. 10.1038/301059a0

Sekimura, T., Fujihashi, Y., & Takeuchi, Y. (2014). A model for population dynamics of the mimetic butterfly Papilio polytes in the Sakishima Islands, Japan. Journal of Theoretical Biology, 361, 133–140. 10.1016/j.jtbi.2014.06.029

Stubblefield, & Seger. (1994). Sexual Dimorphism in the Hymenoptera. In The Differences Between the Sexes.

Trivers, R. L., & Hare, H. (1976). Haplodiploidy and the Evolutiion of the Social Inseicts. 191, 15.

Vane-Wright, R. I. (1971). The systematics of Drusillopsis Oberthür (Satyrinae) and the supposed Amathusiid Bigaena van Eecke (Lepidoptera: Nymphalidae), with some observations on Batesian mimicry. Transactions of the Royal Entomological Society of London, 123(1), 97–123. 10.1111/j.1365-2311.1971.tb00841.x

Virtanen, P., Gommers, R., Oliphant, T. E., Haberland, M., Reddy, T., Cournapeau, D., Burovski, E., Peterson, P., Weckesser, W., Bright, J., van der Walt, S. J., Brett, M., Wilson, J., Millman, K. J., Mayorov, N., Nelson, A. R. J., Jones, E., Kern, R., Larson, E.,…van Mulbregt, P. (2020). SciPy 1.0: Fundamental algorithms for scientific computing in Python. Nature Methods, 17(3), 261–272. 10.1038/s41592-019-0686-2

Waldbauer, G. P. (1970). Mimicry of Hymenopteran Antennaeby Syrphidae. Psyche: A Journal of Entomology, 77(1), 45–49. 10.1155/1970/28967

Waldbauer, G. P., & Sheldon, J. K. (1971). Phenological Relationships of Some Aculeate Hymenoptera, their Dipteran Mimics, and Insectivorous Birds. Evolution, 25(2), 371–382. 10.2307/2406929

Wallace, A. R. (1878). Tropical Nature, and Other Essays. Macmillan and Company.

Waskom, M. L. (2021). seaborn: Statistical data visualization. Journal of Open Source Software, 6(60), 3021. 10.21105/joss.03021

Werren, J. H. (1987). Labile Sex Ratios in Wasps and Bees. BioScience, 37(7), 498–506. 10.2307/1310422

Willadsen, P. C. (2022). Aculeate Hymenopterans as Aposematic and Mimetic Models. Frontiers in Ecology and Evolution, 10, 827319. 10.3389/fevo.2022.827319

Williams, P. (2007). The distribution of bumblebee colour patterns worldwide: Possible significance for thermoregulation, crypsis, and warning mimicry: BUMBLEBEE COLOUR-PATTERN GROUPS. Biological Journal of the Linnean Society, 92(1), 97–118. 10.1111/j.1095-8312.2007.00878.x

Wilson, J. S., Jahner, J. P., Forister, M. L., Sheehan, E. S., Williams, K. A., & Pitts, J. P. (2015). North American velvet ants form one of the world’s largest known Mullerian mimicry complexes. Current Biology, 25(16), R704–R706. 10.1016/j.cub.2015.06.053

Wilson, J. S., Williams, K. A., Forister, M. L., von Dohlen, C. D., & Pitts, J. P. (2012). Repeated evolution in overlapping mimicry rings among North American velvet ants. Nature Communications, 3(1), 1272. 10.1038/ncomms2275

Yamauchi, A. (1993). A population dynamic model of Batesian mimicry. Researches on Population Ecology, 35(2), 295–315. 10.1007/BF02513602

